# Crystal structures of peanut lectin in the presence of synthetic β-N- and β-S-galactosides disclose evidences for the recognition of different glycomimetic ligands

**DOI:** 10.1101/2020.06.20.162875

**Authors:** Alejandro J. Cagnoni, Emiliano D. Primo, Sebastián Klinke, María E. Cano, Walter Giordano, Karina V. Mariño, José Kovensky, Fernando A. Goldbaum, María Laura Uhrig, Lisandro H. Otero

## Abstract

Carbohydrate−lectin interactions are involved in important cellular recognition processes, including viral and bacterial infections, inflammation, and tumor metastasis. Hence, the structural studies of lectin-synthetic glycan complexes are essential for understanding the lectin recognition processes and the further design of promising chemotherapeutics that interfere with sugar-lectin interactions.

Plant lectins are excellent models for the study of the molecular recognition process. Among them, peanut lectin (PNA) is highly relevant in the glycobiology field, because of its specificity for β-galactosides, showing high affinity towards the Thomsen-Friedenreich (TF) antigen, a well-known tumor-associated carbohydrate antigen. Given this specificity, PNA is one of the most frequently used molecular probes for the recognition of tumor cell-surface O-glycans. Thus, it has been extensively used in glycobiology for inhibition studies with a variety of β-galactoside and β-lactoside ligands. Herein, crystal structures of PNA are reported in complex with six novel synthetic hydrolytically stable β-N- and β-S-galactosides. These complexes, along with computational simulations, disclosed key molecular binding interactions of the different sugars to PNA at the atomic level, revealing the role of specific water molecules in the protein–ligand recognition. Furthermore, binding affinity studies measured by isothermal titration calorimetry showed dissociation constant values in the micromolar range, as well as a positive glycoside cluster effect in terms of affinity in the case of the divalent compounds. Taken together, this work provides qualitative structural rationale for the upcoming synthesis of optimized glycoclusters, designed for the study of lectin-mediated biological processes. The understanding of the recognition of β-N- and β-S-galactosides with PNA represents a benchmark in protein-carbohydrate interactions since they are novel synthetic ligands not belonging to the family of O-linked glycosides.

## 1. Introduction

Carbohydrate–protein interactions regulate a myriad of important biological processes, including pathogen recognition, cell adhesion, cell differentiation and apoptosis, glycoprotein synthesis and folding (Varki *et al.*, 2017). Consequently, the study of these interactions and their modulation has far-reaching implications in biology, biotechnology, and drug design (Ernst & Magnani, 2009; Tamburrini *et al.*, 2020). Lectins are sugar-binding proteins responsible for deciphering the information encoded by glycans present in glycoproteins, glycolipids or glycosaminoglycans (Ambrosi *et al.*, 2005; Lis & Sharon 1998). Carbohydrate recognition domains (CRD) present in lectins specifically bind to carbohydrate structures in a reversible manner, mainly through a complex network of hydrogen bonds and hydrophobic interactions, and thus, lectin-glycan lattices are formed, which may cause agglutination of cells (Rabinovich *et al.*, 2007; Dennis & Brewer, 2013).

In contrast to the catalytic site of most enzymes, the ligand binding grooves of lectins are usually shallower; hence, the interactions between monosaccharides and lectins are generally weaker than those observed in enzyme-substrate recognition (Varki *et al.*, 2017). This apparent setback is overcome in natural systems by a multivalent display of sugar residues at the cell surface, which leads to the so-called “cluster glycoside effect” (Lee & Lee, 1995, 2000), where multivalent interactions between lectin and ligands bearing numerous copies of the same carbohydrate group are present (Chabre & Roy, 2010; Sleiman *et al.*, 2015).

Plant lectins provide excellent models for the study of carbohydrate recognition (Sleiman *et al.*, 2015; Varki *et al.*, 2017). Moreover, the carbohydrate binding specificity of plant lectins has made them an active subject of research due to their potential application as diagnostic biomarkers. Indeed, specific plant lectins are being exploited as biological tools to identify altered cell surface glycosylation on pathological conditions, such as inflammation or cancer (Peumans & van Damme, 1998; Sleiman *et al.*, 2015).

The *Arachis hypogaea* lectin (peanut agglutinin, PNA) is one of the most thoroughly studied plant lectins (Banerjee *et al.*, 1994; Lotan *et al.*, 1975). PNA has shown preferential binding towards galactose at the monosaccharide level, although it exhibits a higher affinity towards lactose and, more interestingly, to the Thomsen-Friedenreich (TF) antigen, Gal β(1→3)GalNAc (Ryder *et al.*, 1992), a well-known tumor-associated carbohydrate antigen (Christiansen *et al.*, 2014; Cagnoni *et al.*, 2016). Furthermore, PNA has been shown to be mitogenic for human colonic epithelium and HT29 colorectal cancer cells, thus promoting cell proliferation and tumor growth (Ryder *et al.*, 1992). Consequently, consumption of food rich in PNA, such as peanuts, mushrooms, jackfruit, and others, may be of concern (Belur *et al.*, 2019). More recently, PNA-based approaches have been explored for the efficient and specific detection of the TF-antigen in human samples (Nakase *et al.*, 2017; Kumagai *et al.*, 2019).

PNA has been widely used as a model lectin to structurally characterize the effects of a variety of β-galactoside and β-lactoside ligands (Banerjee *et al.*, 1994; Ravishankar *et al.*, 1997, 1999, 2001; Kundhavai Natchiar *et al.*, 2004, 2006; Goel *et al.*, 2005). However, most of the PNA ligands reported to date correspond to O-linked carbohydrates, which are expected to have limited half-life in biological media since they are sensitive to glycosidase-mediated hydrolysis. Conversely, amino (N*-*) and thio (S*-*) glycosides are more resistant towards enzymatic and acidic hydrolysis, and therefore, present a better bioavailability under physiological conditions (Cagnoni *et al.*, 2011; Cagnoni *et al.*, 2014; Cano *et al.*, 2017; Wang *et al.*, 2018). For this purpose, we used a set of previously synthesized glycoclusters, namely NGS, diNGS, diNGT, STG, STGD, and diSTGD (Fig. 1). These ligands possess β-linked galactose residues and have been proved to be PNA ligands showing different binding affinities (Cagnoni *et al.*, 2014; Cano *et al.*, 2017).

**Figure 1.**
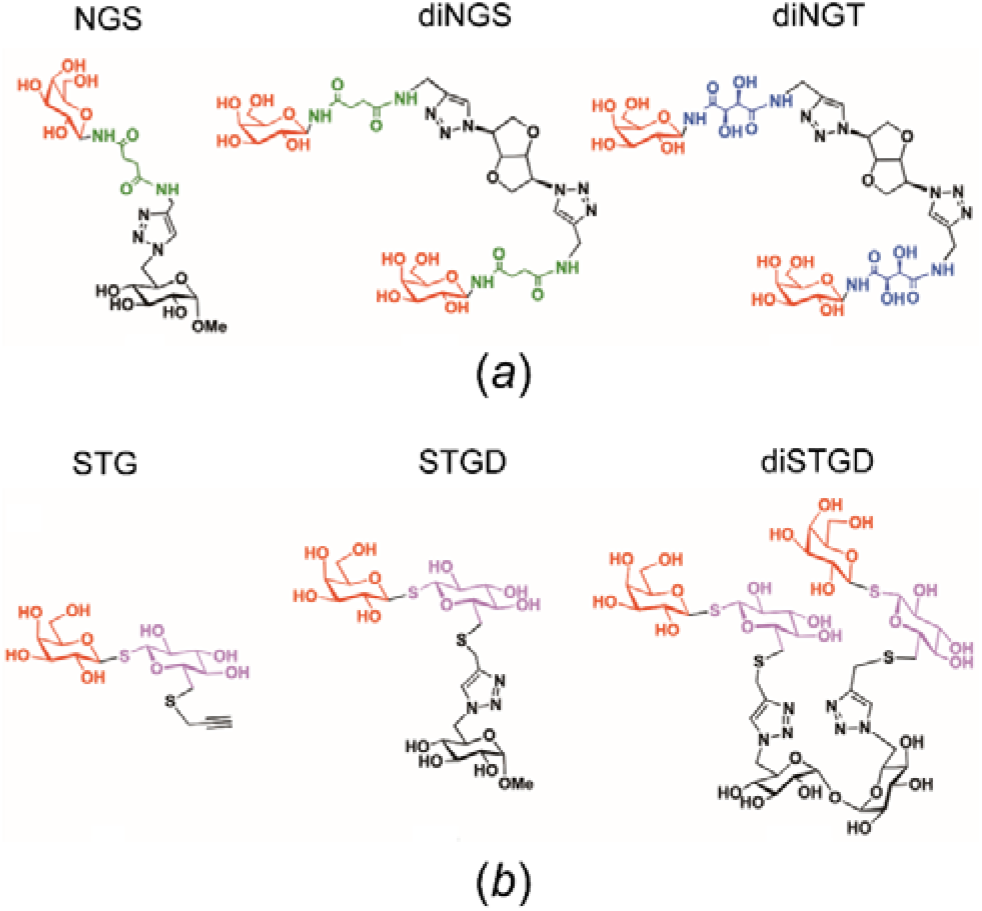
Chemical representations of the synthetic glycoclusters studied in this work: (*a*) N*-*linked glycoclusters NGS, diNGS, diNGT. β-galactoside moieties are depicted in red, succinimidyl linkers in green, tartaramidyl linkers in blue, and triazole rings and sugar scaffolds in black. (*b*) S*-*linked glycoclusters STG, STGD and diSTGD. β-galactoside moieties are depicted in red, β-thioglucopyranosides in purple and propargyl groups, triazole rings and sugar scaffolds in black.

In this work, we present the crystal structures and molecular dynamics studies of PNA in complex with these synthetic glycan ligands and compare them with the PNA-lactose complex previously reported. In addition, we report the isothermal titration calorimetric study of the interaction of the synthetic ligands with PNA. Altogether, the findings herein provided allow a detailed characterization of the PNA-carbohydrate interactions from a thermodynamic and structural point of view, including the role of water molecules.

## 2. Materials and Methods

### 2.1. Reagents

Mature *Arachis hypogaea* lectin (peanut agglutinin, PNA), composed of 236 residues, was purchased from Sigma-Aldrich (L0881, lyophilized powder, affinity purified, agglutination activity <0.1 μg ml^−1^). Compounds NGS, diNGS, diNGT, STG, STGD, diSTGD (Fig. 1) were synthesized as previously described (Cagnoni *et al.*, 2014; Cano *et al.*, 2017).

### 2.2. Crystallization

Initial crystallization conditions for PNA were screened at room temperature by the sitting-drop vapor diffusion method using a Honeybee-963 robot (Digilab, Marlborough, MA, USA) and commercial screens from Hampton Research (Aliso Viejo, CA, USA) and Jena Bioscience (Jena, Germany). Droplets consisted of 350 nl of protein (at 5 mg ml^−1^ in 10 mM Tris-HCl, 25 mM sodium chloride, pH 7.7) and 350 nl of crystallization solution and were set up in 96-well Greiner 609120 plates (Monroe, NC). A total of 10 conditions out of the 576 tested yielded preliminary plate-shaped crystals after a few days of equilibration. The best crystals were eventually grown by the hanging-drop vapor diffusion method by mixing 2 μl of the protein and 2 μl of a precipitation solution consisting of 15% (*w/v*) PEG 8000, 0.1 M sodium citrate, 50 - 125 mM ammonium sulfate, in 24-well Hampton Research VDX plates, and reached a maximum size of 0.30 × 0.15 × 0.05 mm^3^. Ligands were introduced by overnight soaking of the crystals in mother liquor added with 50 mM of the respective compounds. Samples were then cryoprotected in mother liquor added with 20% (*v/v*) 2-methyl-2,4-pentanediol (MPD) and flash-cooled in liquid nitrogen in Hampton Research loops.

### 2.3. Structure resolution, model building, and refinement

Native X-ray diffraction data were collected at 100 K at the PROXIMA-1 and PROXIMA-2A protein crystallography beamlines at Synchrotron SOLEIL (France) using the MXCuBE application (Gabadinho *et al.*, 2010) and then processed with XDS (Kabsch, 2010) and Aimless (Evans & Murshudov, 2013), leaving 5% of the reflection apart for cross-validation. The complex structures were solved by the molecular replacement method with Phaser (McCoy *et al.*, 2007) using the PNA structure as a search model (PDB code 2PEL, Banerjee *et al.*, 1996). The structures were refined with Buster (Bricogne *et al.*, 2011) and manually built with Coot (Emsley *et al.*, 2010). Ligand coordinates were generated with HyperChem^TM^ Professional 7.51, Hypercube Inc. (USA). Missing ligand atoms in the electron density maps were set to zero occupancy. The final models were validated with MolProbity (Chen *et al.*, 2010) and the validation module implemented in Coot (Emsley *et al.*, 2010). The program LigPlot+ (Laskowski *et al.*, 2011) was used for protein-ligand interaction analysis. Table 1 presents detailed information on the data collection parameters and processing statistics.

**Table 1.**
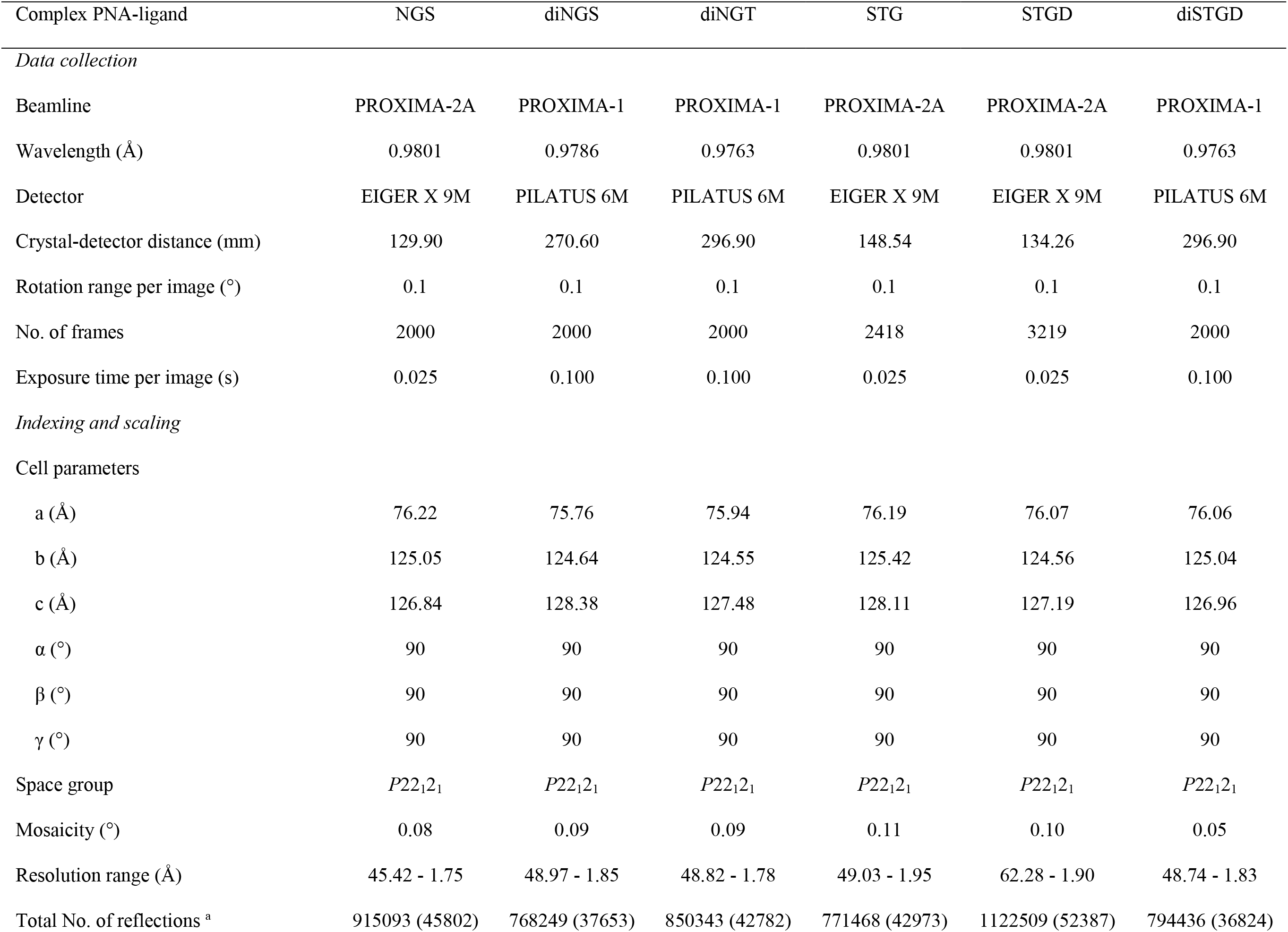
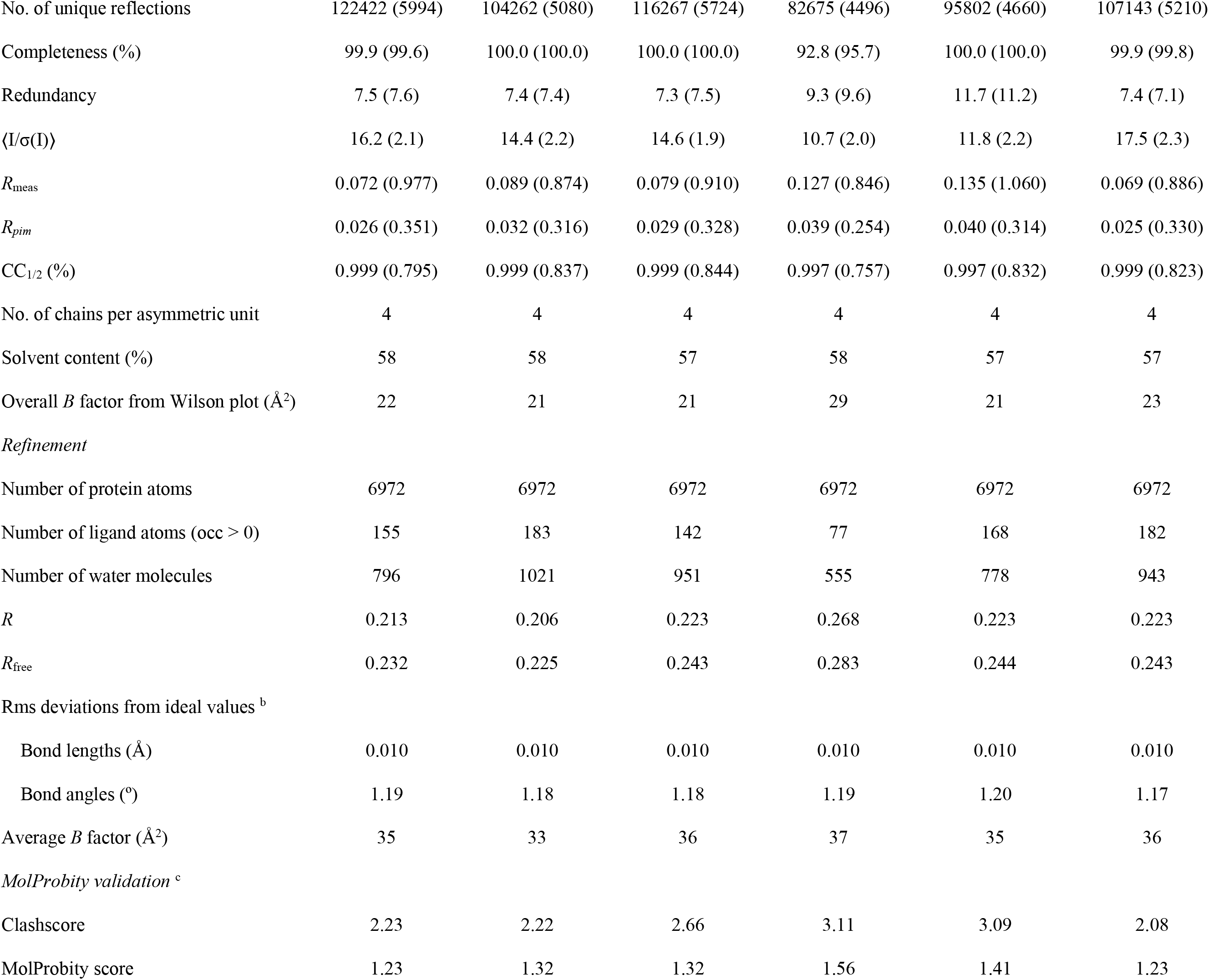
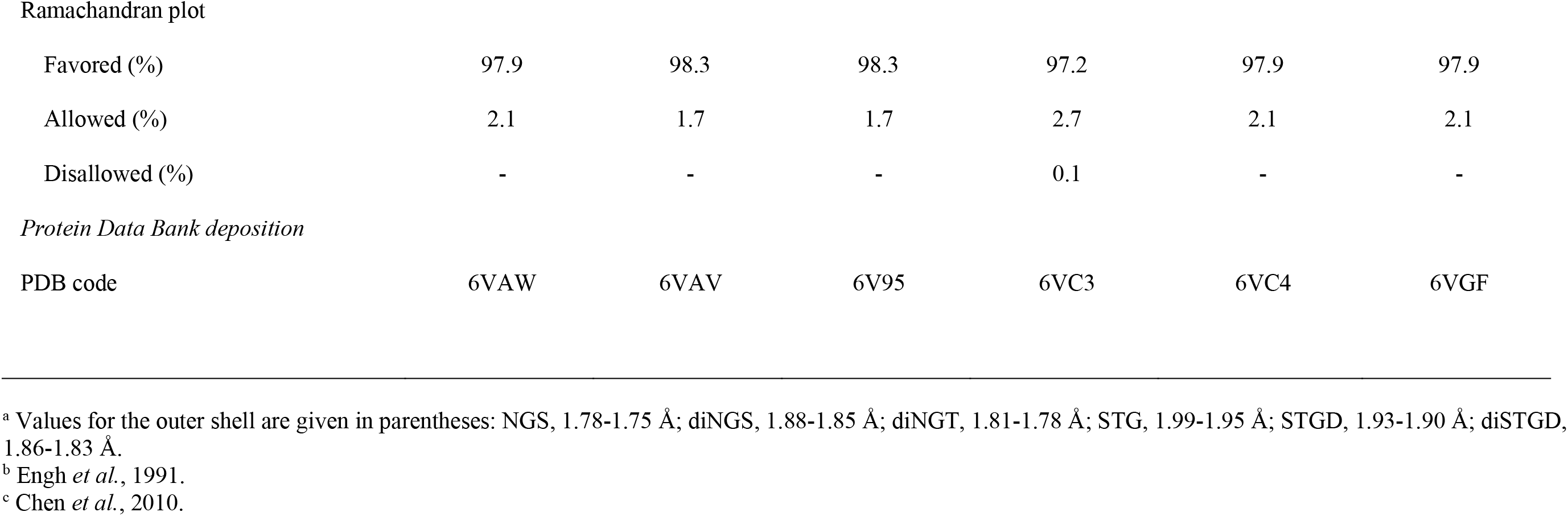
X-ray diffraction data collection and refinement statistics

| Complex PNA-ligand | NGS | diNGS | diNGT | STG | STGD | diSTGD |
| --- | --- | --- | --- | --- | --- | --- |
| *Data collection* |  |  |  |  |  |  |
| Beamline | PROXIMA-2A | PROXIMA-1 | PROXIMA-1 | PROXIMA-2A | PROXIMA-2A | PROXIMA-1 |
| Wavelength (Å) | 0.9801 | 0.9786 | 0.9763 | 0.9801 | 0.9801 | 0.9763 |
| Detector | EIGER X 9M | PILATUS 6M | PILATUS 6M | EIGER X 9M | EIGER X 9M | PILATUS 6M |
| Crystal-detector distance (mm) | 129.90 | 270.60 | 296.90 | 148.54 | 134.26 | 296.90 |
| Rotation range per image (°) | 0.1 | 0.1 | 0.1 | 0.1 | 0.1 | 0.1 |
| No. of frames | 2000 | 2000 | 2000 | 2418 | 3219 | 2000 |
| Exposure time per image (s) | 0.025 | 0.100 | 0.100 | 0.025 | 0.025 | 0.100 |
| *Indexing and scaling* |  |  |  |  |  |  |
| Cell parameters |  |  |  |  |  |  |
| a (Å) | 76.22 | 75.76 | 75.94 | 76.19 | 76.07 | 76.06 |
| b (Å) | 125.05 | 124.64 | 124.55 | 125.42 | 124.56 | 125.04 |
| c (Å) | 126.84 | 128.38 | 127.48 | 128.11 | 127.19 | 126.96 |
| $\alpha$ (°) | 90 | 90 | 90 | 90 | 90 | 90 |
| $\beta$ (°) | 90 | 90 | 90 | 90 | 90 | 90 |
| $\gamma$ (°) | 90 | 90 | 90 | 90 | 90 | 90 |
| Space group | $P22_12_1$ | $P22_12_1$ | $P22_12_1$ | $P22_12_1$ | $P22_12_1$ | $P22_12_1$ |
| Mosaicity (°) | 0.08 | 0.09 | 0.09 | 0.11 | 0.10 | 0.05 |
| Resolution range (Å) | 45.42 - 1.75 | 48.97 - 1.85 | 48.82 - 1.78 | 49.03 - 1.95 | 62.28 - 1.90 | 48.74 - 1.83 |
| Total No. of reflections <sup>a</sup> | 915093 (45802) | 768249 (37653) | 850343 (42782) | 771468 (42973) | 1122509 (52387) | 794436 (36824) |
| No. of unique reflections | 122422 (5994) | 104262 (5080) | 116267 (5724) | 82675 (4496) | 95802 (4660) | 107143 (5210) |
| Completeness (%) | 99.9 (99.6) | 100.0 (100.0) | 100.0 (100.0) | 92.8 (95.7) | 100.0 (100.0) | 99.9 (99.8) |
| Redundancy | 7.5 (7.6) | 7.4 (7.4) | 7.3 (7.5) | 9.3 (9.6) | 11.7 (11.2) | 7.4 (7.1) |
| $\langle I/\sigma(I) \rangle$ | 16.2 (2.1) | 14.4 (2.2) | 14.6 (1.9) | 10.7 (2.0) | 11.8 (2.2) | 17.5 (2.3) |
| $R_{\text{meas}}$ | 0.072 (0.977) | 0.089 (0.874) | 0.079 (0.910) | 0.127 (0.846) | 0.135 (1.060) | 0.069 (0.886) |
| $R_{\text{pim}}$ | 0.026 (0.351) | 0.032 (0.316) | 0.029 (0.328) | 0.039 (0.254) | 0.040 (0.314) | 0.025 (0.330) |
| CC<sub>1/2</sub> (%) | 0.999 (0.795) | 0.999 (0.837) | 0.999 (0.844) | 0.997 (0.757) | 0.997 (0.832) | 0.999 (0.823) |
| No. of chains per asymmetric unit | 4 | 4 | 4 | 4 | 4 | 4 |
| Solvent content (%) | 58 | 58 | 57 | 58 | 57 | 57 |
| Overall $B$ factor from Wilson plot ($\text{\AA}^2$) | 22 | 21 | 21 | 29 | 21 | 23 |
| *Refinement* |  |  |  |  |  |  |
| Number of protein atoms | 6972 | 6972 | 6972 | 6972 | 6972 | 6972 |
| Number of ligand atoms (occ > 0) | 155 | 183 | 142 | 77 | 168 | 182 |
| Number of water molecules | 796 | 1021 | 951 | 555 | 778 | 943 |
| $R$ | 0.213 | 0.206 | 0.223 | 0.268 | 0.223 | 0.223 |
| $R_{\text{free}}$ | 0.232 | 0.225 | 0.243 | 0.283 | 0.244 | 0.243 |
| Rms deviations from ideal values <sup>b</sup> |  |  |  |  |  |  |
| Bond lengths ($\text{\AA}$) | 0.010 | 0.010 | 0.010 | 0.010 | 0.010 | 0.010 |
| Bond angles ($^\circ$) | 1.19 | 1.18 | 1.18 | 1.19 | 1.20 | 1.17 |
| Average $B$ factor ($\text{\AA}^2$) | 35 | 33 | 36 | 37 | 35 | 36 |
| *MolProbity validation* <sup>c</sup> |  |  |  |  |  |  |
| Clashscore | 2.23 | 2.22 | 2.66 | 3.11 | 3.09 | 2.08 |
| MolProbity score | 1.23 | 1.32 | 1.32 | 1.56 | 1.41 | 1.23 |
| Ramachandran plot |  |  |  |  |  |  |
| Favored (%) | 97.9 | 98.3 | 98.3 | 97.2 | 97.9 | 97.9 |
| Allowed (%) | 2.1 | 1.7 | 1.7 | 2.7 | 2.1 | 2.1 |
| Disallowed (%) | - | - | - | 0.1 | - | - |
| Protein Data Bank deposition |  |  |  |  |  |  |
| PDB code | 6VAW | 6VAV | 6V95 | 6VC3 | 6VC4 | 6VGF |

<sup>a</sup> Values for the outer shell are given in parentheses: NGS, 1.78-1.75 Å; diNGS, 1.88-1.85 Å; diNGT, 1.81-1.78 Å; STG, 1.99-1.95 Å; STGD, 1.93-1.90 Å; diSTGD, 1.86-1.83 Å.
<sup>b</sup> Engh *et al.*, 1991.
<sup>c</sup> Chen *et al.*, 2010.

### 2.4. Ligand docking calculations

The publicly available crystal structure of PNA (PDB code 2PEL) was edited for docking calculations using AutoDock Tools 1.5.6. Polar hydrogens and partial charges were added explicitly, whereas the other hydrogen atoms were automatically added by the program. The three-dimensional carbohydrate structures of the glycan ligands were obtained with HyperChem^™^ Professional 7.51 (Hypercube Inc., USA). Ligand structures were optimized with Avogadro 2.0.7 (Hanwell *et al.*, 2012). The edited structure of PNA was used for the docking procedure with AutoDock Vina 1.2 (Trott & Olson, 2010). The most stable conformers of the glycan compounds were manually docked into the carbohydrate-binding sites of PNA by superimposing the terminal Gal residue with that of the crystallographic coordinates. The docking protocol was initially set to rigid conditions with a size of the dock grid of 16 × 16 × 16 Å, which encompasses the binding site for the carbohydrate ligands. Exhaustiveness was initially set to 10 with all other parameters set on default values, and then it was increased to 100 for the final dockings. The Autogrid 4 program present in Autodock 4.2 generated grids of probe atom interaction energies and electrostatic potential. A grid spacing of 0.375 Å was used for the local searches. For each calculation, 100 docking runs were performed using a population of 250 individuals and an energy evaluation number of 3 × 10^6^. The top-ranked complexes, sorted by binding energy values, were visually inspected for good stereochemical geometry and docking and further used as starting conformations for Molecular Dynamics (MD) studies.

### 2.5. Molecular dynamics studies

MD studies were performed with the Amber18 computational simulation package (Case *et al.*, 2018). In all cases, PNA complexes were solvated with explicit three-site point charge modeled (TIP3P) water molecules in an octahedral box, localizing the box limits 10 Å away from the protein surface. MD simulations were performed at 1 atm and 300 K, maintained with the Berendsen barostat and thermostat (Berendsen *et al.*, 1984; van Gunsteren & Berendsen, 1990), using periodic boundary conditions and Ewald sums (grid spacing of 1 Å) for treating long-range electrostatic interactions with a 10 Å cut-off for computing direct interactions. The SHAKE algorithm was applied to all hydrogen-containing bonds, allowing employment of a 2 fs step for the integration of Newton’s equations. Amber ff99SBildn and GLYCAM_06j-1 force field parameters (Hornak *et al.*, 2006) were used for PNA residues and glycans, respectively. The equilibration protocol involved a minimization of the initial structure, followed by a 400 ps constant volume MD run heating the system slowly to 300 K. Finally, a 0.8 ns MD run at constant pressure was performed to achieve proper density. MD runs of 500 ns for all complexes were performed. Frames were saved at 1 ps intervals.

### 2.6. Isothermal Titration Calorimetry (ITC)

All isothermal titration calorimetry experiments were performed by using a NanoITC (TA Instruments). Needle and cell concentrations, injection volumes, and time intervals between injections were varied to obtain sufficient heat production per injection to allow good peak integration, and sufficient time between injections to allow the return to equilibrium (Cagnoni *et al.*, 2014). A typical titration involved 20 injections at 300 s intervals of 2.5 μl aliquots of a 2.5 mM ligand solution into the sample cell (volume 200 μl) containing 50 μM of PNA. The solutions were prepared by dissolving the ligand in water at 298 K. The titration cell was continuously stirred at 300 rpm. The heats of dilution of the ligands in the buffer were subtracted from the titration data. Fitting was performed using the Nano Analyze software to determine the binding stoichiometry (*n*), association constant (*K*_a_), and the enthalpy change (ΔH).

### 2.7. Graphical representation

Molecular structures and their electron densities were represented using PyMOL Molecular Graphics System 1.8 (Schrödinger, USA). Docking poses generated by AutoDock Vina were directly loaded into PyMOL through the PyMOL AutoDock/Vina Plugin for visualization (Seeliger & de Groot, 2010). MD results were visualized with the VMD software 1.9.1 (Humphrey *et al.*, 1996).

## 3. Results and discussion

### 3.1. Crystallographic studies

The ligands used in this work were synthesized as described (Cagnoni *et al.*, 2014; Cano *et al.*, 2017). They can be structurally classified in two distinct subgroups: the first one including compounds NGS, diNGS, and diNGT, where the β-galactose moieties are N-linked through amide functional groups (Fig. 1*a*), and the second one, comprising compounds STG, STGD and diSTGD, where the β-galactoside residues are S*-*linked by thioglycosidic bonds (Fig. 1*b*).

The crystallographic structures of PNA in complex with these six novel synthetic ligands were solved at high resolution (Fig. 2). All crystals belong to the orthorhombic space group *P*22_1_2_1_ and contain four polypeptide chains in the asymmetric unit assembling a homotetramer, as previously described (Banerjee *et al.*, 1994, 1996). In all structures, the electron density maps are continuous, except for the last 19 C-terminal residues in all chains. X-ray data-collection, refinement and stereochemical quality statistics are shown in Table 1.

**Figure 2.**
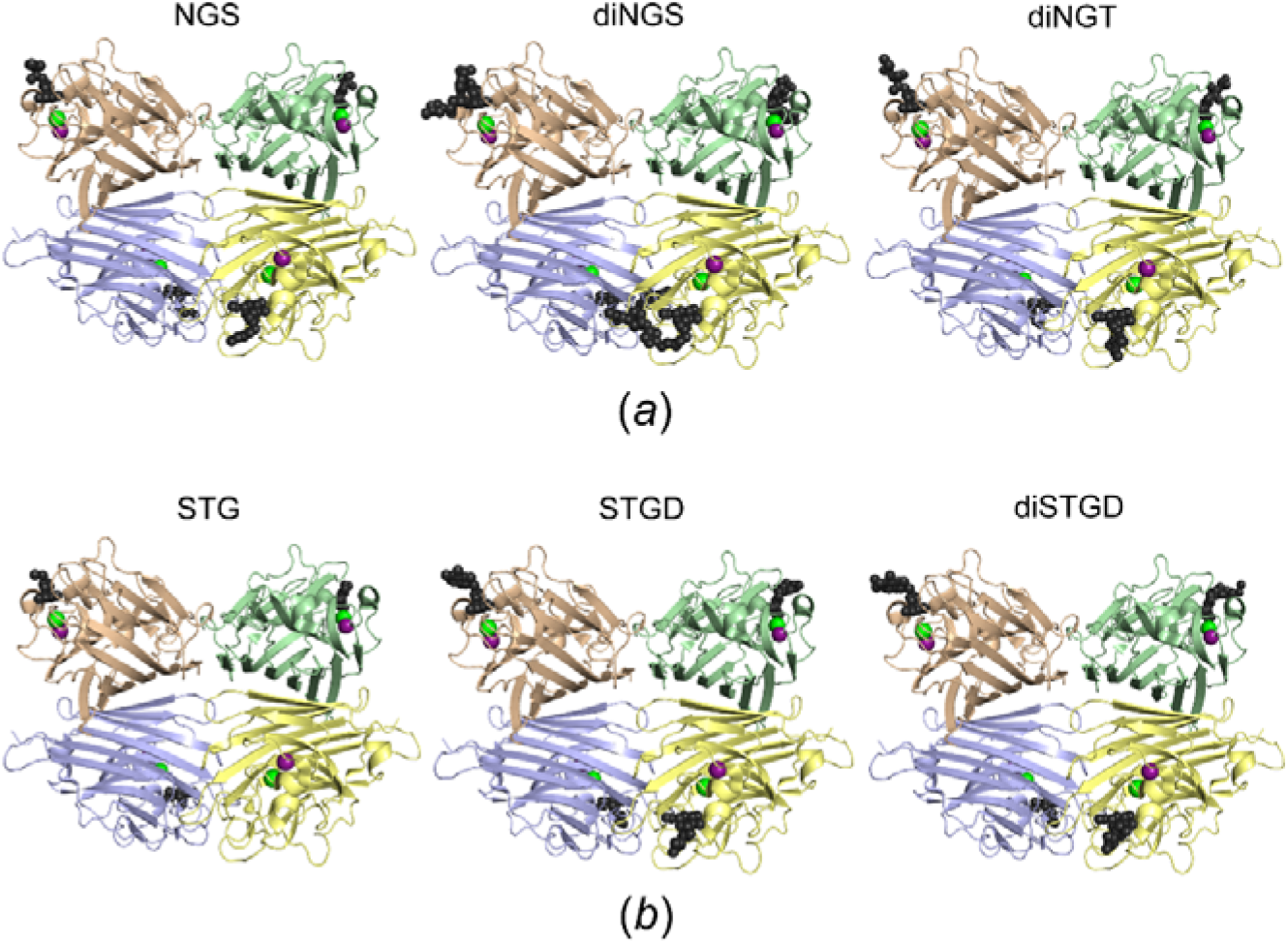
Crystal structures. (*a*) PNA in complex with the N-linked glycoclusters NGS, diNGS, and diNGT. (*b*) PNA in complex with the S-linked glycoclusters STG, STGD, and diSTGD. The four PNA subunits observed in the quaternary assembly are denoted in cartoon representation in different colors. The glycocluster molecules bound to the subunits are represented in black sticks. Only the ligand portions defined by the electron density maps are shown. Magenta and green balls represent Mn^2+^ and Ca^2+^ ions, respectively.

#### 3.1.1. Overall structural features

In all protein-ligand complexes reported in this work, the tertiary and quaternary structures remain largely unaffected in comparison with the crystal structure of PNA in its apo form (Banerjee *et al.*, 1994) and complexed with lactose (Banerjee *et al.*, 1996) with C^α^-r.m.s.d. values between individual chains ranging from 0.11 to 0.22 Å.

Each subunit of PNA complexed with the glycoclusters present the characteristic jelly-roll lectin fold, which consists of a nearly flat six-stranded β-sheet, a curved seven-stranded β-sheet, a small five-stranded β-sheet (linking the two larger β-sheets), and a number of loops of differing lengths and conformations, as reported (Banerjee *et al.*, 1994, 1996; Ravishankar *et al.*, 1999). The quaternary arrangement is mainly stabilized by contacts between the flat six-stranded β-sheets from chains A-D and chains B-C.

The sugar-binding site is shaped by four loops and involves residues Asn41, Asp80, Asp83, Ile101, Gly104, Tyr125, Ser126, Asn127, Ser128, Glu129, Tyr130, Ser211, Leu212, and Gly213, which directly or indirectly interact with the glycan ligand *via* water-mediated contacts, as mentioned in detail below (Banerjee *et al.*, 1996; Ravishankar *et al.*, 2001; Kundhavai Natchair *et al.*, 2006). A calcium ion and a manganese ion are structurally bound near the sugar-binding site in all PNA-ligand complexes (Fig. 2), as previously reported (Banerjee *et al.*, 1994, 1996; Ravishankar *et al.*, 2001; Kundhavai Natchair *et al.*, 2006). The sugar-binding sites of the four subunits in all crystal complexes are occupied with their respective ligands, with the exception of the PNA-STG complex, chain C (Fig. 2). However, in all crystal complexes, a large fraction of each of the ligand molecules does not present electron density. It is well known that amide and thioether linkages, as well as triazole rings, are highly stable at physiologic pH (*i.e*. in the crystallization conditions employed) (Greenber *et al.*, 2002; Tron *et al.*, 2008). Furthermore, N- and S-glycosidic bonds have been shown to be resistant to enzymatic hydrolysis (Cagnoni *et al.*, 2014; Cano *et al.*, 2017; Sulzenbacher *et al.*, 1997; Driguez *et al.*, 1994, 2001). Consequently, this observation is attributed to the fact that a significant portion of ligand molecules lie outside the sugar-binding site, and are thus totally disordered, as explained in detail below.

#### 3.1.2. Binding of the galactose moiety in the different synthetic glycoclusters

In all protein-ligand structures, the galactose moiety shows clearly defined electron density along with the bound water molecules around it at the sugar-binding pocket (Fig. 3, and Supplementary Figs. S1 and S2). A structural comparison among the different protein complexes indicates an unperturbable position of this sugar ring with no significant changes in the binding pocket residues (Supplementary Fig. S3). The only remarkable divergence is the location of the Asp80 residue, whose side chain adopts different conformations not only among the six ligand-protein complexes but also between the different subunits from the same complex (Supplementary Fig. S3).

**Figure 3.**
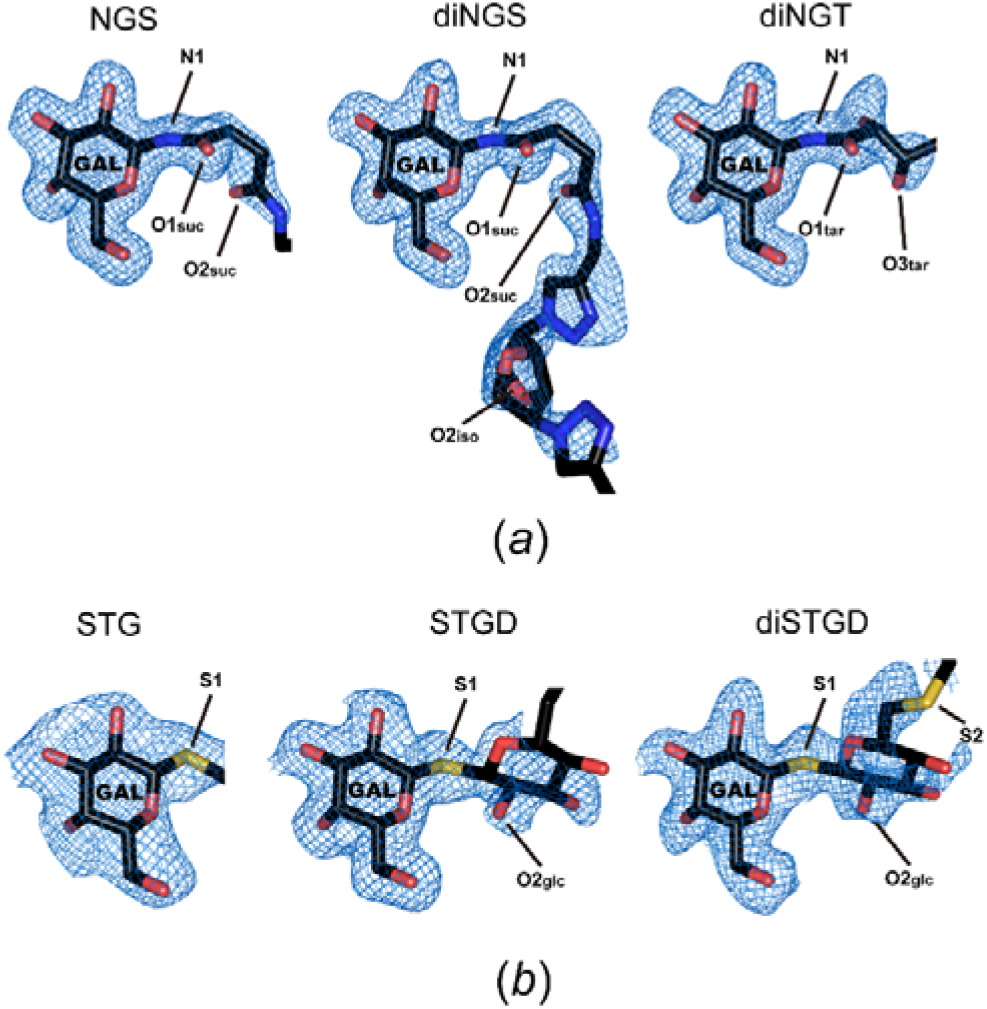
Final *2mF*o–*DF*c electron density maps around the bound synthetic glycoclusters. (*a*) N-linked glycoclusters NGS, diNGS, and diNGT. (*b*) S-linked glycoclusters STG, STGD, and diSTGD. The orientation is similar for all ligands. The maps (blue mesh) are contoured at the 1.0 σ level. The ligands are shown as sticks with carbon atoms in black, oxygen atoms in red, nitrogen atoms in blue, and sulfur atoms in yellow. Only ligand portions defined by the electron density maps are shown. The galactose moiety is labelled for clarity. Oxygen atoms from the succinimidyl and tartaramidyl carbonyl group adjacent to the amide linkage are labeled as O1_suc_ and O1_tar_, respectively. Oxygen atoms from the succinimidyl and tartaramidyl carbonyl group distal to the amide linkage are indicated as O2_suc_ and O3_tar_, respectively. The oxygen atom from the symmetric isomannide scaffold is labeled as O2_iso_. The oxygen atom from the thioglucose is labeled as O2_glc_. The nitrogen and sulfur atoms from the amide and thioglucoside linkages are labeled as N1, S1, and S2, respectively. The maps were generated in PHENIX (Adams *et al.*, 2010).

The protein-galactose atomic interactions are nearly identical to those observed in the PNA-β-galactosides complexes previously reported (Fig. 4) (Banerjee *et al.*, 1994, 1996; Ravishankar *et al.*, 1997, 1999, 2001; Kundhavai Natchiar *et al.*, 2006). Briefly, O2_gal_ interacts with two ordered water molecules, W1 and W2 (the latter was not observed in the PNA-STG complex). W1 is stabilized by the amide group of Gly104 and a hydrogen-bonded network involving water molecules W3, W4, W5 and W6 coordinated by the backbones of Ile101 (carbonyl group) and Leu212 (amide group), along with the side chains of Asn41, Glu129 and Tyr130. On the other hand, W2 is found at hydrogen bond distance from the Glu129 carboxylate, and two extra water molecules (W7 and W8) are coordinated by the backbones of Ser126 (carbonyl group) and Ser128 (amide group), and the hydroxyl group and the O^γ^ atom of Tyr125 and Ser128, respectively. Moreover, O3_gal_ interacts with the carboxylate of Asp83, the side chain of Asn127 and the amide group of Gly104. Furthermore, O4_gal_ is coordinated by the carboxylate group of Asp83 along with the side chain of Ser211, and O5_gal_ is mainly stabilized by the side chain of Ser211. Additionally, O6_gal_ is in close contact with the carboxylate group of Asp80, but as noted above, this interaction is not always present due to the different orientations of this residue. Additionally, the side chain of Tyr125 stacks against the galactose ring at the primary binding site (CH/π bond) as previously reported (Asensio *et al.*, 2013). The direct and water-mediated protein-ligand interactions in the different crystallographic structures are summarized in Tables 2 and 3. Taken all together, our results show that, in all complexes, the β-galactoside residues present a highly conserved conformation in the ligand binding pocket, and thus, the lectin-glycan interactions are in accordance with those previously reported (Banerjee *et al.*, 1996; Ravishankar *et al.*, 1997, 1999, 2001; Kundhavai Natchiar *et al.*, 2004, 2006; Goel *et al.*, 2005). Thus, as expected, the β-galactoside moieties present in the synthetic N- and S-linked glycoclusters represent the main recognition elements for lectin binding.

**Table 2.** Direct interactions between PNA and ligands at the sugar-binding site

| Interacting atoms |  | Ligands |  |  |  |  |  |  |
| --- | --- | --- | --- | --- | --- | --- | --- | --- |
| Protein | Ligand | N-linked glycoclusters |  |  | S-linked glycoclusters |  |  | Lactose |
|  |  | NGS | diNGS | diNGT | STG | STGD | diSTGD |  |
| Asp83 OD1 | O3<sub>gal</sub> | 2.65 | 2.60 | 2.70 | 2.77 | 2.58 | 2.63 | 2.55 |
| Asn127 ND2 | O3<sub>gal</sub> | 2.85 | 2.93 | 2.93 | 2.83 | 3.10 | 2.90 | 3.00 |
| Gly104 N | O3<sub>gal</sub> | 3.13 | 3.10 | 2.98 | 3.10 | 3.03 | 3.00 | 2.98 |
| Asp83 OD2 | O4<sub>gal</sub> | 2.65 | 2.68 | 2.73 | 2.87 | 2.68 | 2.68 | 2.60 |
| Ser211 OG | O4<sub>gal</sub> | 2.65 | 2.70 | 2.63 | 2.77 | 2.70 | 2.73 | 2.75 |
| Ser211 OG | O5<sub>gal</sub> | 3.33 | 3.30 | 3.28 | 3.40 | 3.20 | 3.20 | 3.18 |
| Asp80 OD2 | O6<sub>gal</sub> | 3.18 | 2.73 | 3.40 | 3.20 | 2.85 | 2.95 | 3.28 |
| Ser211 OG | O3<sub>glc</sub>* | - | - | - | - | - | - | 3.33 |
| Gly213 N | O3<sub>glc</sub>* | - | - | - | - | - | - | 3.18 |

The distances (Å) are averaged over all subunits
Polar interactions within hydrogen-bonding distance (cutoff < 3.4 Å) are considered.
\*only observed in the PNA-lactose complex.

**Table 3.** Water-mediated interactions between PNA and ligands at the sugar-binding site

| Water | Interacting atoms |  |  | Ligands |  |  |  |  |  |  |
| --- | --- | --- | --- | --- | --- | --- | --- | --- | --- | --- |
|  | Protein | Waters | Ligand | N-linked glycoclusters |  |  | S-linked glycoclusters |  |  | Lactose |
|  |  |  |  | NGS | diNGS | diNGT | STG | STGD | diSTGD |  |
| W1 | Gly104 N | W3, W6 | O2<sub>gal</sub> | X | X | X | X | X | X | X |
| W2 | Glu129 OE2 | W7 | O2<sub>gal</sub> | X | X | X | X | X | X | X |
| W3 | Ile101 O | W1, W4 | - | X | X | X | X | X | X | X |
| W4 | - | W3, W5 | N1, O1 | X | X | X | - | - | - | X |
| W5 | Asn41 ND2, Leu212 N | W4 | - | X | X | X | X | X | X | X |
| W6 | Glu129 OE2, Tyr130 OH | W1 | - | X | X | X | X | X | X | X |
| W7 | Tyr125 OH | W2, W8 | - | X | X | X | - | X | X | - |
| W8 | Ser126 O, Ser128 OG, Ser128N | W7* | - | X | X | X | X | X | X | X |
| W9 | Ser211 OG, Gly213 N | - | O2<sub>suc</sub>, O3<sub>tar</sub>, S1, O2<sub>glc</sub><sup>#</sup> | X | X | X | X | X | X | - |
| W10 | Tyr125 OH | - | O1<sub>suc</sub>, O1<sub>tar</sub> | X | X | X | - | - | - | - |
| W11 | Asp80 OD2 | - | O2<sub>iso</sub> | - | X | - | - | - | - | - |
| W12 | - | W7 | S2 | - | - | - | - | - | X | - |

X= Presence of the water molecule at the PNA sugar-binding site.
Polar interactions within hydrogen-bonding distance (cutoff < 3.4 Å) are considered.
\*not observed in the PNA-STG and PNA-lactose complexes.
<sup>#</sup> not observed in the PNA-STG complex.

**Figure 4.**
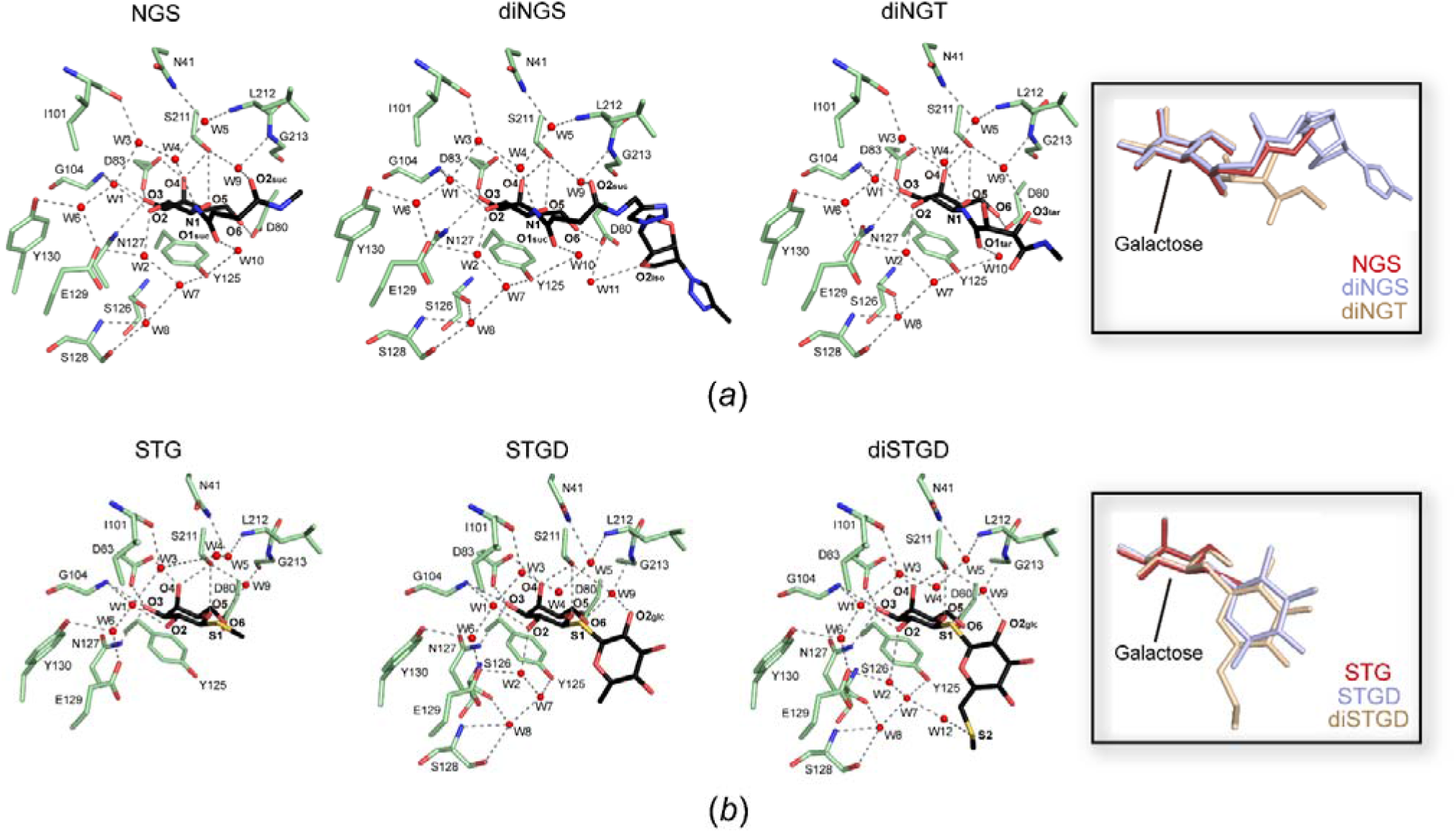
Interactions of the synthetic glycoclusters with the PNA sugar-binding pocket. (*a*) PNA complexed with the N-linked glycoclusters NGS, diNGS, and diNGT. (*b*) PNA complexed with the S-linked glycoclusters STG, STGD, and diSTGD. In all lectin–ligand complexes similar orientations are presented. The ligands are shown as sticks with carbon atoms in black, oxygen atoms in red, nitrogen atoms in blue, and sulfur atoms in yellow. Only the ligand portions defined by the electron density maps are shown. The most relevant residues involved in the interactions are depicted as sticks with carbon atoms in green, oxygen in red, and nitrogen in blue. Water molecules in the active site are shown as red spheres. Galactose oxygen atoms are labeled as O2, O3, O4, O5, and O6 in all ligands. Oxygen atoms from the succinimidyl and tartaramidyl carbonyl group adjacent to the amide linkage are labeled as O1_suc_ and O1_tar_, respectively. Oxygen atoms from the succinimidyl and tartaramidyl carbonyl group distal to the amide linkage are labeled as O2_suc_ and O3_tar_, respectively. The oxygen atom from the symmetric isomannide scaffold is labeled as O2_iso_. The oxygen atom from the thioglucose is labeled as O2_glc_. The nitrogen and sulfur atoms from the amide and thioglucoside linkages are labelled as N1, S1, and S2, respectively. Polar interactions within hydrogen-bonding distance (cutoff < 3.4 Å) are shown as dashed lines. The insets highlight the structural contrasts of the different N*-*linked and S*-*linked glycocluster conformations buried into the PNA sugar-binding pocket. The galactose moiety is indicated for clarity in each case.

#### 3.1.3. Binding of the N-linked glycoclusters beyond the galactose moiety

As mentioned above, while the β-N-galactosyl residue present in NGS, diNGS and diNGT could be clearly detected in the CRD domain, the distal moieties are not supported by the electron density maps (Fig. 3 and Supplementary Fig. S1). As expected, these ligand atoms are facing the solvent and do not contact the protein (Cano *et al.*, 2017). Moreover, the simultaneous presence of two β-galactoside recognition elements from divalent ligands in two distinct binding sites of a PNA tetramer was not observed, which could be attributed to the short linkers of the glycoclusters. However, the second β-galactoside residue contributes to the increased lectin affinity observed for the divalent ligands diNGS and diNGT by ITC experiments (see below).

Nevertheless, in addition to the galactose moiety described above, the amide linkage and the succinimidyl (NGS and diNGS) or tartaramidyl (diNGT) moieties are well defined into their electron density maps, since additional stabilizing contacts take place around these portions of the ligands.

In the three ligands, the N1 atom from the amide linkage makes a hydrogen bond with the water molecule W4 from the constellation of contacts interacting with the O2_gal_ atom (Fig. 4*a* and Table 3). Moreover, there is an interaction between the oxygen atoms from the succinimidyl and tartaramidyl carbonyl groups adjacent to the amide linkage (O1_suc_ and O1_tar_, respectively) with the side chain of Tyr125 *via* the water molecule W10 (Fig. 4*a* and Table 3).

In the glycan ligands encompassing a succinimidyl chain (NGS and diNGS), the oxygen atom from the carbonyl group distal to the amide linkage (O2_suc_) is coordinated by the water molecule W9 that is stabilized by the side chain of Ser211 and the amide group of Gly213. Interestingly, in the diNGT ligand, the same coordination network is found to be interacting with the tartaramidyl *R*-configured hydroxyl oxygen atom (O3_tar_) distal to the amide linkage (Fig. 4*a* and Table 3). Notably, although tartaramidyl and succinimidyl linkers are essentially different due to the presence or absence of the hydroxyl groups, in both cases, an oxygen atom occupies the same position, suggesting that the water-bridged interaction with W9 is essential for linker stabilization.

Compared to β-galactoside disaccharides, the N-linked NGS, diNGS and diNGT ligands, bearing a linear functionalized chain bound to the β-Gal group, exhibit a higher flexibility. Notably, in all PNA-ligand complexes with the synthetic N-linked glycoclusters, oxygen atoms present at a γ- or δ-carbon atom from the anomeric position are consistently disposed towards Ser211, Leu212 and Gly213, interacting with the W9 water molecule and, in turn, with PNA residues. The observation is in accordance with our previous hypothesis, assuming that the presence of a conveniently disposed hydroxyl group in the spacer, as in diNGT, could emulate the role of the 3’-OH group in lactose (O3_glc_) in the PNA-lactose complex (see details below), as was shown by molecular docking studies (Cano *et al.*, 2017). The results obtained here confirm that the linker carbonyl oxygen atoms are also involved in these interactions.

Moreover, in the PNA-diNGS complex, the electron density map for the ligand is exceptionally continuous up to the symmetric isomannide scaffold (Fig. 3 and Supplementary Fig. S1). However, the only relevant interaction detected is a water-bridged hydrogen bond (W11) between an isomannide oxygen atom (O2_iso_) and the side chain of Asp80 (Fig. 4*a* and Table 3). Nevertheless, the observed interaction reflects the fact that not only the recognition element but also the carbohydrate scaffold may present additional binding interactions with the lectin residues or the water-network at the ligand binding pocket, enhancing the ligand affinity. Historically, carbohydrate-based scaffolds have not been extensively investigated compared to dendrimers or polymers (Kiessling *et al.*, 2006; Arsiwala *et al.*, 2019). However, sugar scaffolds have previously shown promising properties for the design of multivalent glycoclusters, based on improved hydrophilicity and pharmacokinetics, as compared to peptidic, aromatic, or polymeric scaffolds (Gouin *et al.*, 2007).

#### 3.1.4. Binding of the S-linked glycoclusters beyond the galactose moiety

As observed also for the N-linked glycoclusters, a considerable fraction of the molecules of the three synthetic thioglycoclusters STG, STGD and diSTGD is missing in the electron density maps of the complex structures. Nevertheless, the regions adjacent to the galactose moiety such as the thioglucoside linkage and the nearby β-thioglucose residue (not observed in STG) are well defined (Fig. 3 and Supplementary Fig. S1).

In this regard, in the three protein-ligand complexes, the S1 sulfur atom linking the galactose and β-thioglucose moieties is coordinated by the ordered water molecule W9, instead of W4 observed in the amide linkage binding (Fig. 4*b* and Table 3). Although the presence of W4 is detected, the contact distance (>3.7 Å) is not suitable for hydrogen bond interaction with the thioglucoside linkage (Fig. 4*b* and Table 3). Additionally, it should be noted that sulfur hydrogen bonds are usually weaker than O-mediated interactions (Biswal, 2015). Furthermore, an extra atomic interaction can be appreciated between W9 and the O2_glc_ oxygen atom from the thioglucopyranose ring in the STGD and diSTGD complexes. As a result, the β-thioglucose ring seems to be stabilized, in contrast to STG, which does not present electron density (Fig. 3 and Supplementary Fig. S1). Interestingly, the O2_glc_ atom in the thioglycoclusters STGD and diSTGD occupies a nearly analogous position to that of O2_suc_ from the distal carbonyl group in the NGS and diNGS succinimidyl chains (Fig. 4 and Supplementary Fig. S3). These results suggest that, besides the tightly bound β-galactose residue, the spacer on N-linked glycoclusters, or the β-glucopyranose ring on the S-linked glycans, also adopt a relatively conserved conformation, allowing a water-bridge interaction with residues Ser211 and Gly213, being W9 the key water molecule for the ligand stabilization.

Furthermore, in the PNA-diSTGD complex, further interactions are observed in the ligand binding beyond the β-thioglucose ring. Here, the second thioglucosidic bond (S2) is found within hydrogen-bonding distance from a water molecule (W12) that is coordinated by W7 from the W2 water-mediated interaction network with O2_gal_ (Fig. 4*b* and Table 3). This additional interaction could be related to the trehalose scaffold, only present on diSTGD, which allows a closer disposition of the β-glucopyranose ring towards PNA.

#### 3.1.5. Structural comparison between the interactions of the synthetic glycoclusters and lactose at the PNA binding site

As previously noted, the tertiary and quaternary structure of PNA in complex with the synthetic glycoclusters is identical as that reported in the complex with lactose (Banerjee *et al.*, 1996). Moreover, the position and binding interaction of the galactopyranose ring is equivalent, where all the direct and water-mediated hydrogen-bonded interactions (except for water W7) along with the non-polar contacts are preserved (Fig. 5*a* and Tables 2 and 3). Regarding the linkage of the different compounds, the O1 oxygen atom from the β-(1→4) glycosidic bond of lactose is found 1.2 Å closer to the protein (from the Ser211 O^γ^ atom as reference) with respect to the N1 atom from the amide linkage, but almost in similar place in comparison with the S1 atom from the β-(1→1) thioglucoside bond (Fig. 5*b*). However, as previously mentioned, the water-bridged interactions observed with the amide linkage of the N-linked glycoclusters remain unalterable in the O-glycosidic linkage from lactose, as illustrated in Figs 4*a* and 5*a*, respectively.

**Figure 5.**
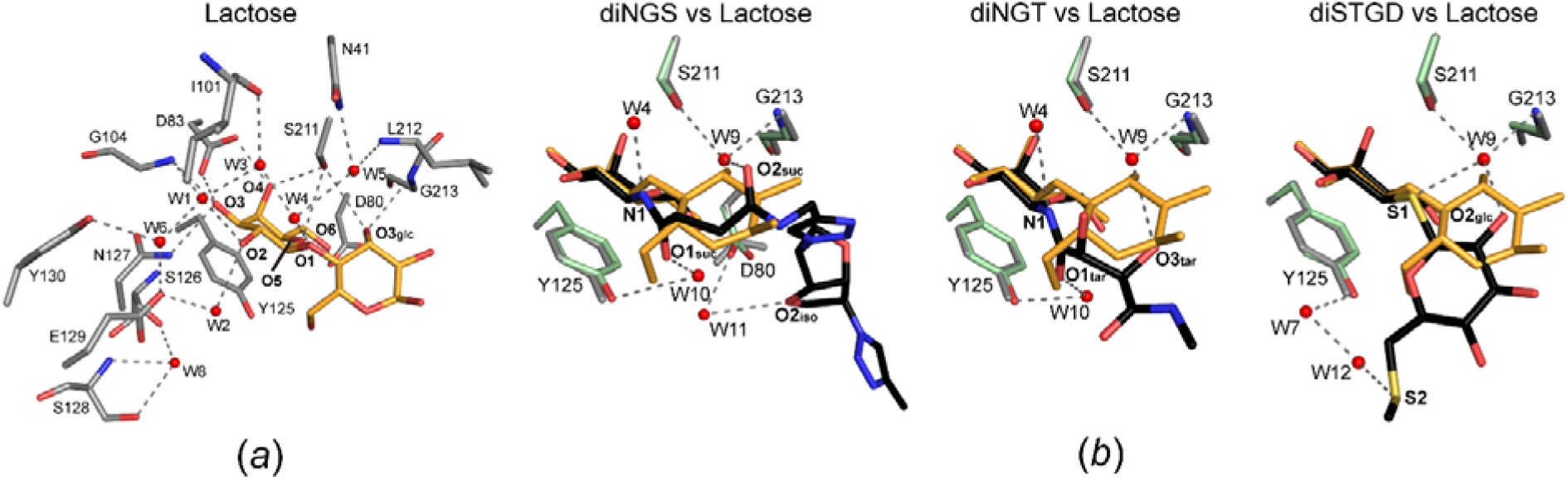
Comparison of the interactions of PNA with lactose and PNA with the divalent synthetic glycoclusters. (*a*) Lactose binding mode as observed in the PNA-lactose crystallographic complex (PDB code: 2PEL). Lactose is shown as sticks with carbon atoms in orange and oxygen atoms in red. The most relevant residues involved in the interactions are depicted as sticks with carbon atoms in gray, oxygen in red, and nitrogen in blue. Galactose oxygen atoms are labeled as O2, O3, O4, O5, and O6. The oxygen atom O3_glc_ from the neighboring glucopyranose ring is marked. (*b*) Structural contrasts between the binding interactions of diNGS, diNGT and diSTGD with respect to lactose at the PNA sugar-binding site. The ligands along with the most relevant residues from the PNA-diNGS, PNA-diNGT and PNA-diSTGD complexes are colored according to Fig. 4. Only the ligand portions defined by the electron density maps are shown. Atoms from the galactose-appended chemical moieties are labeled according to Fig. 4. Lactose and residues from the PNA-lactose complex are depicted as orange and gray sticks, respectively. Crucial water molecules in the active site are shown as red spheres. The most relevant polar interaction differences beyond galactose moiety are stressed as dashed lines in the PNA-diNGS, PNA-diNGT and PNA-diSTGD complexes.

Interestingly, in the lactose complex the glucopyranose ring interacts with PNA by means of hydrogen bonding interactions between the O3_glc_ atom and the side chain of Ser211 and the amide group of Gly213 (Fig. 5*a*). In the synthetic glycoclusters, the water molecule W9 is found nearly in the same location of the lactose O3_glc_ atom, and thus stabilized by the same residues (Fig. 5*b*). Remarkably, as detailed above, W9 plays a crucial role in the interactions of the PNA binding pocket with the succinimidyl or tartaramidyl groups of the N-linked glycoclusters, as well as with the β-thioglucopyranose ring of the S*-*linked glycoclusters. Furthermore, a structural comparison between lactose and the thioglycoclusters (both bearing a glucopyranose moiety) reveals a displacement in the glucose position of around 3 Å (Fig. 5*b*, right). Interestingly, when designing the thioglycoclusters bearing thiodigalactoside (TDG) analogues as recognition elements, we hypothesized that the glucose ring conformation would be well-preserved. However, this is not the actual scenario. This conformational dislocation is probably a consequence of the differences in the linkage of the glucose to the galactose (β(1→4), lactose; β(1→1), thioglycoclusters), which generates a 180° axial rotation of the glucose ring disposition. Moreover, the significantly longer anomeric carbon-sulfur bond, compared to an O-glycosidic linkage, could also explain this difference.

### 3.2. Molecular dynamics of the PNA-synthetic glycocluster complexes

Based on the evidence obtained by the analysis of the crystal structures, molecular dynamics (MD) simulations of apo PNA and in complex with NGS and STG were carried out to provide further insights into the molecular interactions. NGS and STG were selected as the structurally representative members of the distinct groups of N-linked and S-linked synthetic glycan ligands, respectively. The results showed that the evaluated glycans adopt essentially the same conformations in the ligand binding pocket of PNA (Fig. 6*a*) than those observed in the crystallographic complexes (Fig. 4). Concordantly, most of the β-galactoside atoms were in close contact with the lectin residues, showing protein-galactose interactions nearly identical to those observed in the PNA-β-galactosides complexes, while the linker (on NGS) and the β-glucopyranose ring (on STG) are located facing the solvent (Fig. 6*a*).

**Figure 6.**
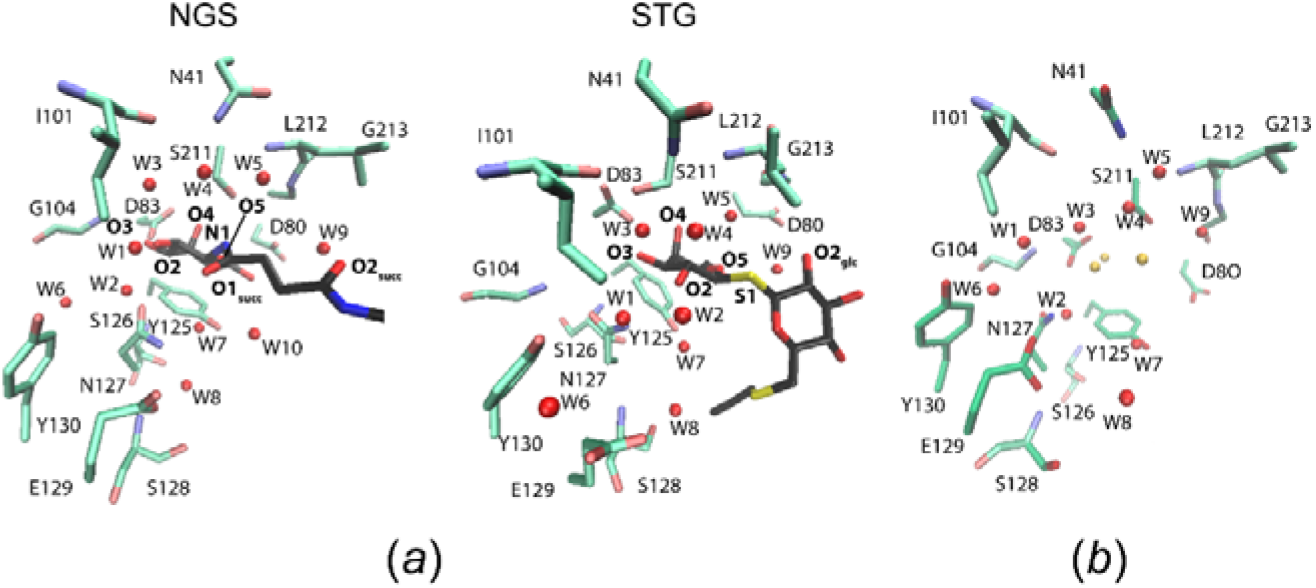
Molecular dynamics simulations at the PNA sugar-binding site. (*a*) PNA complexed with the glycoclusters NGS and STG. (*b*) Hydration in apo PNA. Similar orientations are presented in all cases. Protein residues and ligands are depicted using the same color conventions as in Figure 4. Water molecules from clusters 1 and 2 are colored in yellow and red, respectively.

In addition, on both computational simulations, the water network and the main water-bridged interactions between the ligands and PNA were consistent with the ones observed in the crystallographic studies. Particularly, water molecules W4 and W9, critical for the stabilization of the ligand (in NGS) and the β-glucopyranose ring (in STG), respectively, adopted the same disposition as those observed in the crystallographic complexes (Fig. 6*a*). Remarkably, the water molecules W2, W7, and W8 were also observed in the PNA-STG MD simulation, despite not being clearly defined in the crystal structure (Fig. 4).

Interestingly, in the MD simulation of apo PNA, a highly conserved water-network could be observed (Fig. 6*b*). Compared to the PNA-synthetic ligand complexes, two distinct set of water molecules can be recognized. On the one hand, a group of water molecules (named cluster 1) are closely located to the hydroxyl groups of the galactose moieties, namely O1_gal_, O3_gal_ and O4_gal_. In this sense, MD studies of the solvent adjacent to the protein surface in lectin-ligand complexes have allowed the characterization of regions with higher water occupancy than the bulk solvent, named water sites (Gauto *et al.*, 2013). These sites mimic the interaction between the carbohydrate hydroxyl groups and the lectin, and have been used for improved docking methods for the study of several lectin-glycan interactions (Guardia *et al.*, 2011; Gauto *et al.*, 2011; Modenutti *et al.*, 2015). On the other hand, a second set of specific water molecules (named cluster 2), including W1, W3, W4, W5, W6, W7 and W9, adopt very close dispositions to the ones observed in the crystal structures with the synthetic glycoclusters (Fig. 6*b*). Notably, the water molecule W10, present in the crystallographic complexes of PNA with the N-linked glycoclusters but absent in the coordination with the S-linked synthetic glycans (see Fig. 4), was not observed in the MD simulations of apo PNA. Thus, W10 is coordinated in the sugar-binding site along with the N-linked compound binding.

These results suggest that the PNA sugar-binding site is highly hydrated in the absence of the galactosylated ligands. In the presence of the ligand, water molecules from cluster 1 are replaced by the galactosyl residue; however, water molecules from cluster 2 are preserved in nearly the same positions being key for ligand stabilization.

### 3.3. Binding affinity of the synthetic glycoclusters to PNA by ITC

In order to determine the association constants (*K*_a_) and thermodynamics of the binding of the synthetic glycoconjugates to PNA, Isothermal Titration Calorimetry (ITC) experiments were performed using the same conditions as reported previously (Cagnoni *et al.*, 2014). All glycoclusters were evaluated as ligands for PNA, using lactose and TDG as reference compounds. The ITC plots were analyzed and the independent model, *i.e.* single site with 1:1 stoichiometry, was sufficient to give a good fit (Supplementary Fig. S4). The relevant thermodynamic binding parameters are reported in Table 4.

**Table 4.** Thermodynamic binding parameters of the synthetic ligands towards PNA related to lactose and TDG, used as Reference^a^

| Compound | Valence | Type | $K_d$<br>($\mu\text{M}$) | $K_a$<br>($\text{M}^{-1}$) | $n$ | Rel. Pot. | $\Delta H$<br>(kJ/mol) | -T.$\Delta S$<br>(kJ/mol) | $\Delta G$<br>(kJ/mol) |
| --- | --- | --- | --- | --- | --- | --- | --- | --- | --- |
| Lactose | 1 | Lactoside (reference) | 643.8 | 1,553.5 | 1.05 | 1.00 | -29.84 | 11.59 | -18.25 |
| TDG | 1 | Thiodigalactoside (TDG, reference) | 523.6 | 1,910.0 | 1.01 | 1.23 | -31.89 | 13.16 | -18.73 |
| STG | 1 | Analogue of TDG | 551.9 | 1,812.0 | 0.97 | 1.17 | -30.13 | 11.60 | -18.53 |
| STGD | 1 | Analogue of TDG | 420.3 | 2,379.0 | 1.03 | 1.53 | -40.93 | 21.67 | -19.27 |
| diSTGD | 2 | Analogue of TDG | 160.6 | 6,225.0 | 0.48 | 4.01 | -75.99 | 54.33 | -21.66 |
| NGS | 1 | Galactosylamine (succinimidyl linker) | 1616.8 | 618.5 | 1.02 | 0.40 | -25.49 | 9.280 | -16.21 |
| diNGS | 2 | Galactosylamine (succinimidyl linker) | 698.1 | 1,432.5 | 0.47 | 1.10 | -70.88 | 52.98 | -17.90 |
| diNGT | 2 | Galactosylamine (tartaramidyl linker) | 544.2 | 1,837.5 | 0.51 | 1.18 | -75.34 | 56.82 | -18.52 |

<sup>a</sup>Valence refers to the structural valence of the ligand; $n$ is the stoichiometry of the binding; Rel. pot. is the relative potency of the ligand referred to reference 1 (lactose).

All tested synthetic compounds showed affinity towards PNA, and the isotherms for the titration of PNA against the synthetic glycoconjugates indicated exothermic behavior at 298 K. As a general trend, the S-linked glycoclusters STG, STGD and diSTGD, bearing TDG analogues as recognition elements, showed higher binding affinity than the β-N-galactosides NGS, diNGS and diNGT (Table 4), in accordance with previous results (Cagnoni *et al.*, 2014; Cano *et al.*, 2017). When comparing the S-linked monovalent glycoclusters STG and STGD to lactose, the synthetic glycans exhibited a slight increase in binding affinity, as observed by the higher *K*_a_ values (Table 4). In this sense, the longer and more flexible C-S thioglycosidic bond and the rotation observed for the β-glucopyranose ring could be playing a favorable role enhancing the affinity of the synthetic glycans for PNA, although this is not evident from the observation of the crystal structures (Fig. 4*b*). Moreover, among the S-linked glycoclusters, the divalent compound diSTGD showed the highest affinity among the evaluated ligands. This enhanced affinity could be attributed to two distinct features: i) the additional water-bridged interactions with the S2 sulfur atom observed in the diSTGD complex (see Fig. 4*b*) and ii) the divalent nature of diSTGD and its multivalent effect. This cluster effect has been previously observed by interaction of PNA with a variety of multivalent O-glycosides (Ambrosi *et al.*, 2005; Gouin *et al.*, 2010; Srinivas *et al.*, 2005).

In the N-linked glycoclusters, the β-glucopyranose ring has been replaced by a flexible linear linker. This structural fact provoked a reduction in the binding affinity towards PNA, compared to lactose, TDG and the S-linked glycoclusters. Indeed, the monovalent ligand NGS showed ca. two-fold lower binding affinity for PNA than for lactose. This reduced affinity is partially compensated by a moderate cluster glycoside effect in the divalent ligands diNGS and diNGT. Moreover, the diNGT ligand, bearing a tartaramidyl linker, exhibited a slightly higher *K*_a_ value than the diNGS glycocluster, *i.e*. 1432.5 and 1837.5 M^−1^, respectively. Accordingly, enzyme-linked lectin assays (ELLA) experiments performed on plates coated with a lactose-polystyrene glycopolymer, using horseradish peroxidase-labelled PNA, also showed a slightly positive effect for the diNGT ligand in terms of affinity, attributed to the presence of the hydroxyl groups of the tartaramidyl spacer (Cano *et al.*, 2017). Consistently, the crystal structures evidenced the participation of specific linker oxygen atoms in water-mediated interactions with the lectin.

In all cases, the thermodynamic profiles of the calorimetric studies showed a significant enthalpic contribution, indicating a well-defined ligand binding groove, as well as a negative entropic contribution, resulting in *K*_a_ association constants of an order of 10^3^ M^−1^. These features reveal enthalpically-driven interactions for the PNA-glycoclusters complexes, which are mainly associated with direct hydrogen bonds established between the lectin and the glycan ligands (Sindrewicz *et al.*, 2019). Thus, these results suggest that CH/π stacking and water-bridged interactions, observed in the crystallographic complex structures, may play a secondary role compared to the conserved direct hydrogen bond interactions of the galactose moieties present in all ligands evaluated. The ΔΔH between the ligand presenting the highest affinity towards PNA, namely diSTGD, compared to the lowest one, NGS, was approximately −50 kJ mol^−1^, reflecting the effect of the structural features present in the evaluated ligands. This difference was mostly compensated by the entropic component, as the (-T. ΔΔS) value was around 45 kJ mol^−1^. Thus, the Gibbs energies of binding (ΔG) varied modestly for these compounds (ΔΔG = −5.45 kJ mol^−1^, Table 4). These enthalpy–entropy compensations are a widely observed phenomenon that occurs with the addition of functional groups to the ligand that leads to tighter van der Waals contacts and hydrogen bonds with the protein (giving a more negative ΔH), while at the same time reducing the translational, rotational and conformational entropy of the protein and/or the ligand (Dragan *et al.*, 2017). In these compounds, however, the highest binding enthalpies and Gibbs energies are also associated with the presence of divalent glycoclusters, presenting higher affinity per residue than the corresponding monovalent compounds. Consistently, the entropy values observed for the divalent compounds diSTGD, diNGT and diNGS were ca. two-fold or more than those observed for the monovalent compounds, indicating a necessarily higher order for the system.

## 4. Final remarks

When designing lectin ligands, several features should be considered, including affinity, selectivity, toxicity and pharmacokinetics. In this sense, a better bioavailability and a higher lifetime in biological fluids may be expected for synthetic glycoclusters comprising N- o S-linkages, compared to that of O-glycosides, based on their resistance to enzymatic hydrolysis mediated by galactosidases.

The structural analysis of PNA in complex with isosteric analogues of natural O-linked glycosides contributes to the understanding of the atomic determinants involved in the recognition process in these cases. With this purpose, we have described six crystal structures of PNA bound to novel synthetic hydrolytically-stable β-N- and β-S-galactoside glycoclusters. To our knowledge, these crystal structures represent the PNA-glycan complexes with the highest resolutions reported to date.

The crystallographic structures along with the computational studies confirmed that hydrogen bond interactions, hydrophobic CH/π stackings and multi-water bridges involved in the recognition process of β-galactoside moieties by PNA are conserved with the synthetic glycoclusters in comparison with lactose. Besides the β-galactoside residue, the structures of all synthetic glycoclusters exhibited a conserved water-bridged interaction with Ser211 and Gly213, in which the water molecule W9 notably occupies the same position as O3_glc_ in the PNA-lactose complex. Affinity studies showed that the thiodigalactoside analogs presented higher binding affinity towards PNA than the β-galactosylamides assayed, and that the divalent ligands exhibited a moderate cluster glycoside effect.

This multidisciplinary study based on X-ray crystallography, molecular modelling simulations and affinity binding measurements, provides valuable information for the further design of new generations of high-affinity lectin ligands, resistant to biological degradation by enzymes.

## Acknowledgments

Support for this work from the Agencia Nacional de Promoción Científica y Tecnológica (ANPCyT) under grants PICT 2015-0621 and PICT 2016-1425, the Consejo Nacional de Investigaciones Científicas y Técnicas (CONICET), and the Universidad de Buenos Aires (UBA), Argentina, are gratefully acknowledged. AJC, SK, WG, KVM, FAG, MLU and LHO are research members of CONICET. EDP would like to acknowledge fellowship support from ANPCyT. We are grateful for the access to the PROXIMA 1 and PROXIMA 2A beamlines at the Synchrotron SOLEIL, France.

## Accession numbers

Coordinates and structure factors have been deposited in the Protein Data Bank (http://wwpdb.org/) under the following codes: 6VAW (PNA-NGS), 6VAV (PNA-diNGS), 6V95 (PNA-diNGT), 6VC3 (PNA-STG), 6VC4 (PNA-STGD), and 6VGF (PNA-diSTGD).

