## Supplementary material for "Crystal structures of peanut lectin in the presence of synthetic β-N- and β-S-galactosides disclose evidences for the recognition of different glycomimetic ligands": Suplementary files

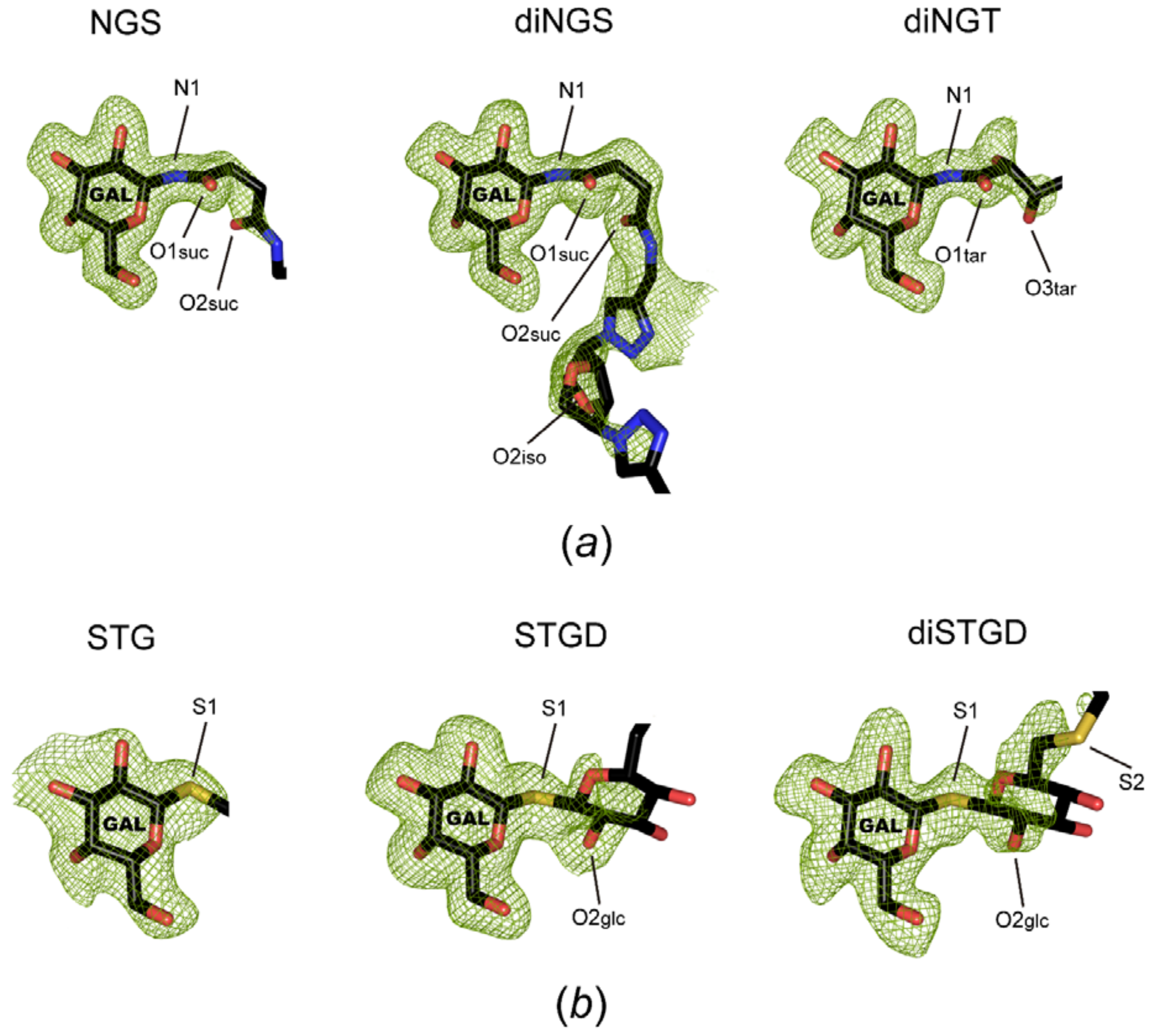


**Figure S1**
Final *mF*o−*DF*c omit difference maps around the bound synthetic glycoclusters. (*a*) N-linked glycoclusters NGS, diNGS, and diNGT. (*b*) S-linked glycoclusters STG, STGD, and diSTGD. Labels, orientations and representations are similar as in Figure 3. Only ligand portions defined by the electron density maps are shown. Omit maps (green mesh) are contoured at 2.5 σ evidencing the presence of the ligands. Omit maps were generated by removing the ligands from the models followed by five cycles of refinement in PHENIX (Adams *et al.*, 2010).

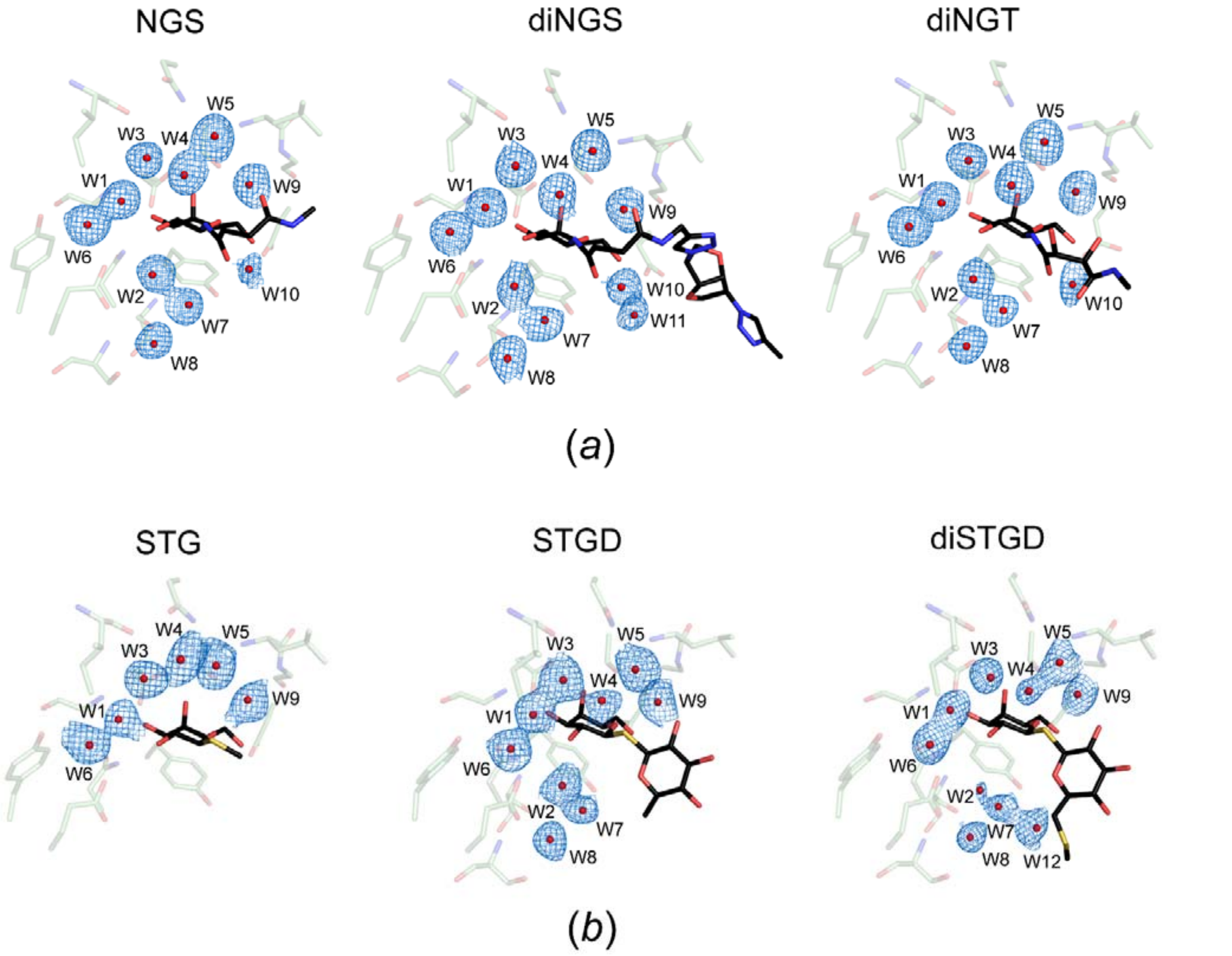


**Figure S2.**

Final *2mF*o–*DF*c electron density maps around the bound water molecules at the sugar-binding pocket of PNA in complex with the different synthetic glycoclusters. (*a*) N-linked glycoclusters NGS, diNGS, and diNGT. (*b*) S-linked glycoclusters STG, STGD, and diSTGD. Similar views with respect to Fig. 4 are shown in all cases. The maps (blue mesh) are contoured at 1.0 σ. Water molecules in the active site are shown as red spheres. The ligands and the most relevant residues involved in the interactions are colored according to Fig. 4. Only ligand portions defined by the electron density maps are shown. The maps were generated in PHENIX (Adams *et al.*, 2010).

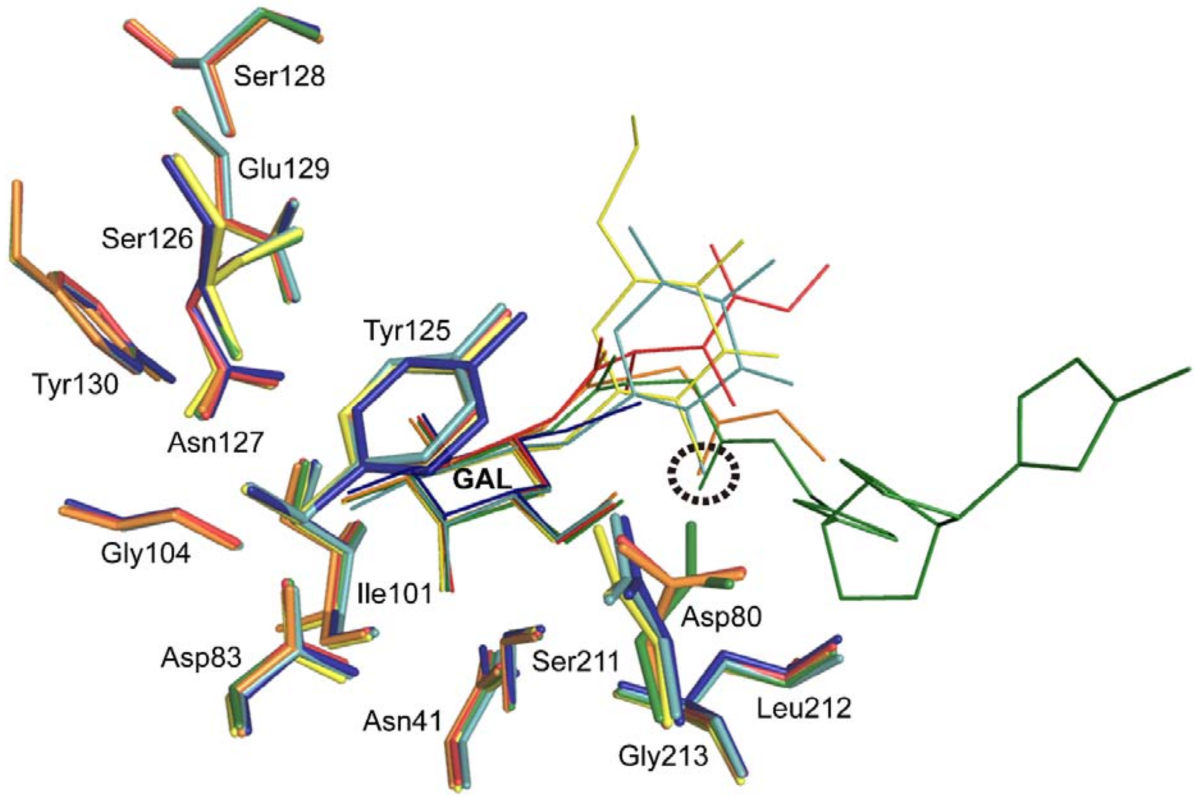


**Figure S3**
Ligand binding structural comparison amongst the different PNA-synthetic glycocluster complexes. The protein-ligand complexes are colored as follows: PNA-NGS, orange; PNA-diNGS, green; PNA-diNGT, red; PNA-STG, blue; PNA-STGD, cyan; and PNA-diSTGD, yellow. The most relevant residues involved in the interactions are depicted as sticks. The ligands are shown as lines and colored according to the complex. Only the ligand portions defined by the electron density maps are shown. With a dashed black circle, the analogous positions between the atoms $O2_{glc}$ from the thioglycoclusters (STGD and diSTGD) and $O2_{suc}$ from the succinimidyl chains (NGS and diNGS) are highlighted. The galactose moiety is labeled for clarity.

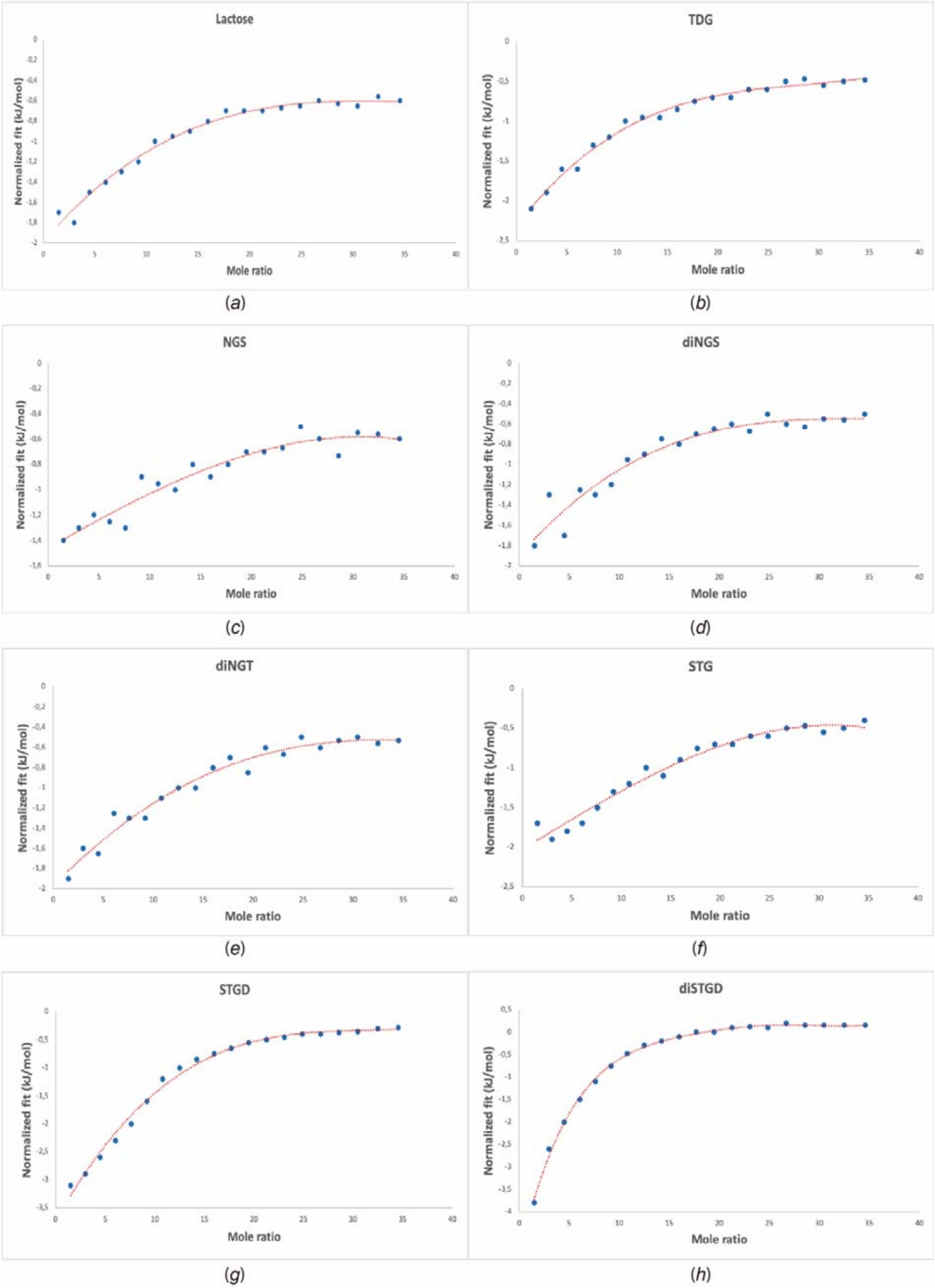


**Figure S4**

Interaction analysis of the synthetic glycoclusters with PNA by ITC. Integrated heats of interaction between PNA and (*a*) lactose, (*b*) TDG, (*c*) NGS, (*d*) diNGS, (*e*) diNGT, (*f*) STG, (*g*) STGD, and (*h*) diSTGD at 298 K. The independent model was implemented using the NanoAnalyze software to obtain the fitting curve for the experimental data.
